# Benchmark of tools for *in silico* prediction of MHC class I and class II genotypes from NGS data

**DOI:** 10.1101/2022.04.28.489842

**Authors:** Arne Claeys, Jasper Staut, Peter Merseburger, Kathleen Marchal, Jimmy Van den Eynden

## Abstract

The Human Leukocyte Antigen (HLA) genes are a group of highly polymorphic genes that are located in the Major Histocompatibility Complex (MHC) region on chromosome 6. The HLA genotype affects the presentability of tumour antigens to the immune system. While knowledge of these genotypes is of utmost importance to study differences in immune responses between cancer patients, gold standard, PCR-derived genotypes are rarely available in large Next Generation Sequencing (NGS) datasets. Therefore, a variety of methods for *in silico* NGS-based HLA genotyping have been developed, bypassing the need to determine these genotypes with separate experiments. However, there is currently no consensus on the best performing tool. Here, we compiled a list of 13 HLA callers and evaluated their accuracy on three different datasets. Based on these results, best-practice guidelines were constructed, and consensus HLA allele predictions were made for DNA and RNA samples from The Cancer Genome Atlas (TCGA).

## Introduction

The human Major Histocompatibility Complex (MHC) is a gene complex located on the p-arm of chromosome 6 that contains two large clusters of genes with antigen processing and presentation functions: the MHC class I and MHC class II regions [1–3].

MHC class I molecules are involved in the presentation of endogenous antigens to cytotoxic T-cells and consist of a heavy chain encoded by one of the Human Leukocyte Antigen Class I genes (*HLA-A, HLA-B* or *HLA-C)*, and a light β_2_ microglobulin chain [4–6]. Their role in tumour immunity has been established for a long time. Indeed, they can present neoantigens, small mutated peptides, to CD8+ T cells, resulting in an immune response and cancer cell death.

The most frequently studied MHC class II genes include *HLA-DPA1, HLA-DPB1, HLA-DQA1, HLA-DQB1, HLA-DRA1* and *HLA-DRB1*. They encode alpha and beta heterodimers that form the MHC class II protein complex. The role of these genes in anti-tumour immunity is emerging [7–9]. MHC class II mediated tumour-immune interaction occurs either via an indirect or a direct mechanism. First, cancer cells can secrete neoantigens that are subsequently taken up and presented on the MHC-II of antigen presenting cells infiltrating the tumour [7,10,11]. Additionally, some tumours express MHC-II themselves and can directly interact with CD4+ T-cells [7,11].

The peptide-binding region of HLA molecules is highly polymorphic and specific HLA alleles determine neoantigen binding and presentation to the immune system. Genotype dependent differences in HLA binding affinity could lead to differential responses to immunotherapy, as illustrated by the association that has been described between MHC-I genotypes (e.g., *HLA-B62*) and survival in immune checkpoint blockade (ICB)-treated advanced melanoma patients [12]. It is currently unclear whether MHC-II genotypes also determine responses to immunotherapy. Such association studies require knowledge of the HLA genotype. PCR methods are currently the gold standard for this genotyping but datasets with PCR-based HLA calls are rarely available [13–15]. HLA genotyping can also be performed on Next Generation Sequencing (NGS) data. A plethora of tools has been developed for this task. *Polysolver* and *Optitype* are often recommended as the best performing tools for MHC-I genotyping [16]. For MHC-II genotyping there is currently no consensus about the best method. Several benchmarks have been performed previously [14,16–23], but these were either not applied to MHC class II or did not include some recently published tools.

In this study, we compiled a list of 13 tools that predict HLA genotypes from NGS data and benchmarked their performance on both the 1000 genomes dataset and on an independent cell line dataset. Subsequently we assessed their performance on 9162 DNA and 9761 RNA sequencing files from The Cancer Genome Atlas (TCGA) by comparing the predicted allele frequencies with reference population allele frequencies. Based on these findings, we give recommendations on which tool to use for a given data type and how the outputs of multiple tools can be combined into a consensus prediction.

## Results

### Selection of 13 HLA genotyping tools with variable computational resource requirements

We identified 22 available HLA genotyping tools from literature (Table 1). Thirteen tools that were free for academic use, applicable on Whole Exome Sequencing (WES), Whole Genome Sequencing (WGS) or RNA-Seq data and ran on Ubuntu 20.04 were included in this study: *arcasHLA, HLA-HD, HLA-VBSeq, HLA*LA, HLAforest, HLAminer, HLAscan, Kourami, Optitype, PHLAT, Polysolver, seq2HLA* and *xHLA (*Table S1). These HLA typing algorithms mainly differ in the method to align reads to the HLA allele reference sequences and in the subsequent allele prioritization approach (Table S2). All 13 tools can make allele predictions for the three MHC class I genes (*HLA-A, HLA-B* and *HLA-C*) and 9 tools support additional calling of the MHC class II genes *HLA-DPA1, HLA-DPB1, HLA-DQA1, HLA-DQB1* and *HLA-DRB1*. Two methods support only a subset of the MHC class II genes: *xHLA* does not support calling *HLA-DPA1* and *HLA-DQA1*, while *PHLAT* does not support *HLA-DPA1* and *HLA-DPB1*. The tools also differ in which data types they support: 6 of them require DNA data, 3 tools require RNA data and the 4 remaining tools support both data types (Table 1).

**Table 1.**
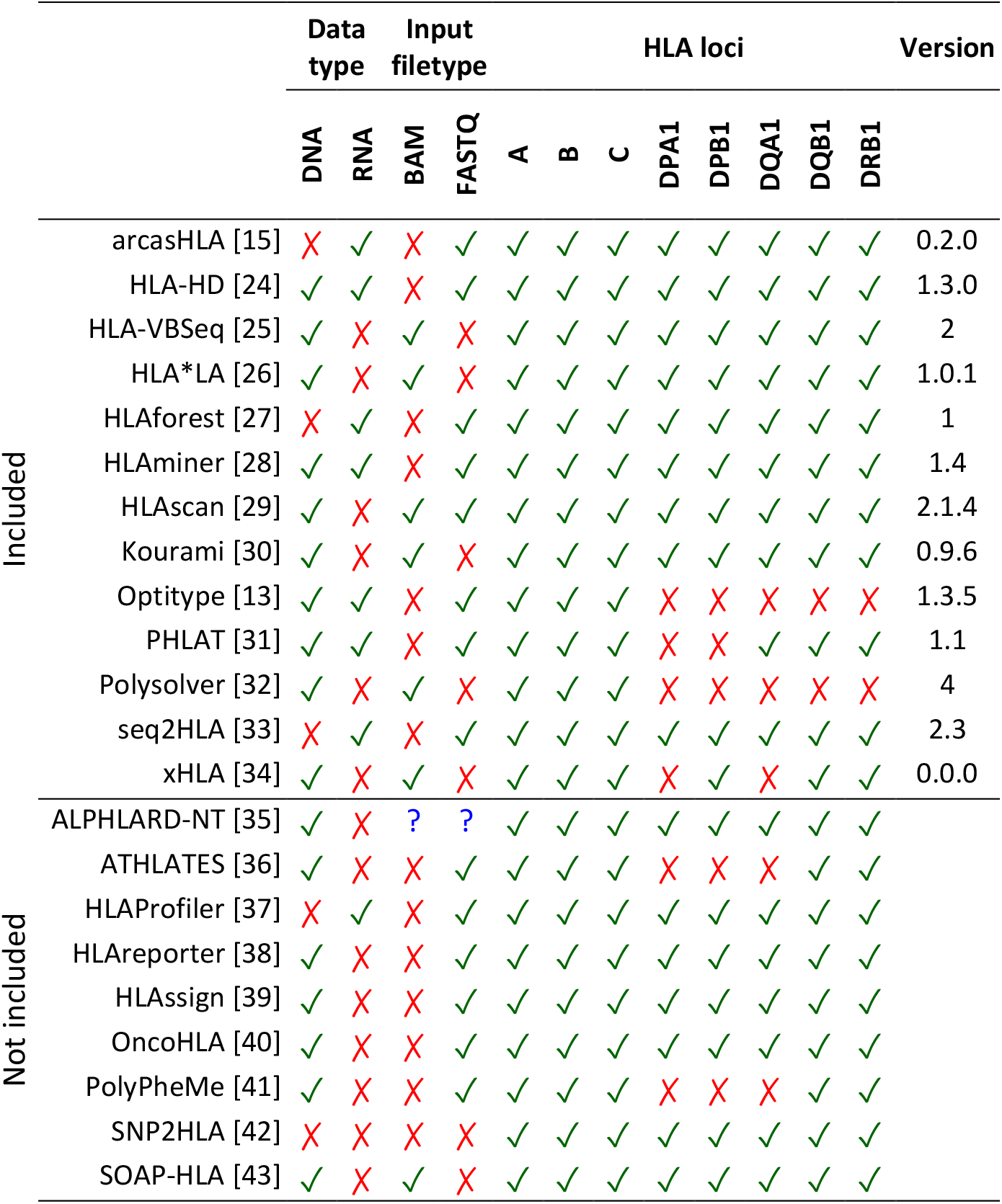
Overview of evaluated tools for HLA genotyping. List of tools for NGS-based HLA genotyping. Checkmarks and crosses indicate which NGS methods (DNA-Seq and/or RNA-Seq) and input file types (FASTQ and/or BAM) are supported and for which genes predictions can be made. The tools in the upper part of the table are benchmarked in this study. Tools in the lower part of the table did not fulfil our inclusion criteria and were not further considered.

Firstly, the computing time and memory usage of the thirteen selected tools were measured on a random subset of 10 DNA and 10 RNA sequencing files from the TCGA project (Fig 1).

**Figure 1.**
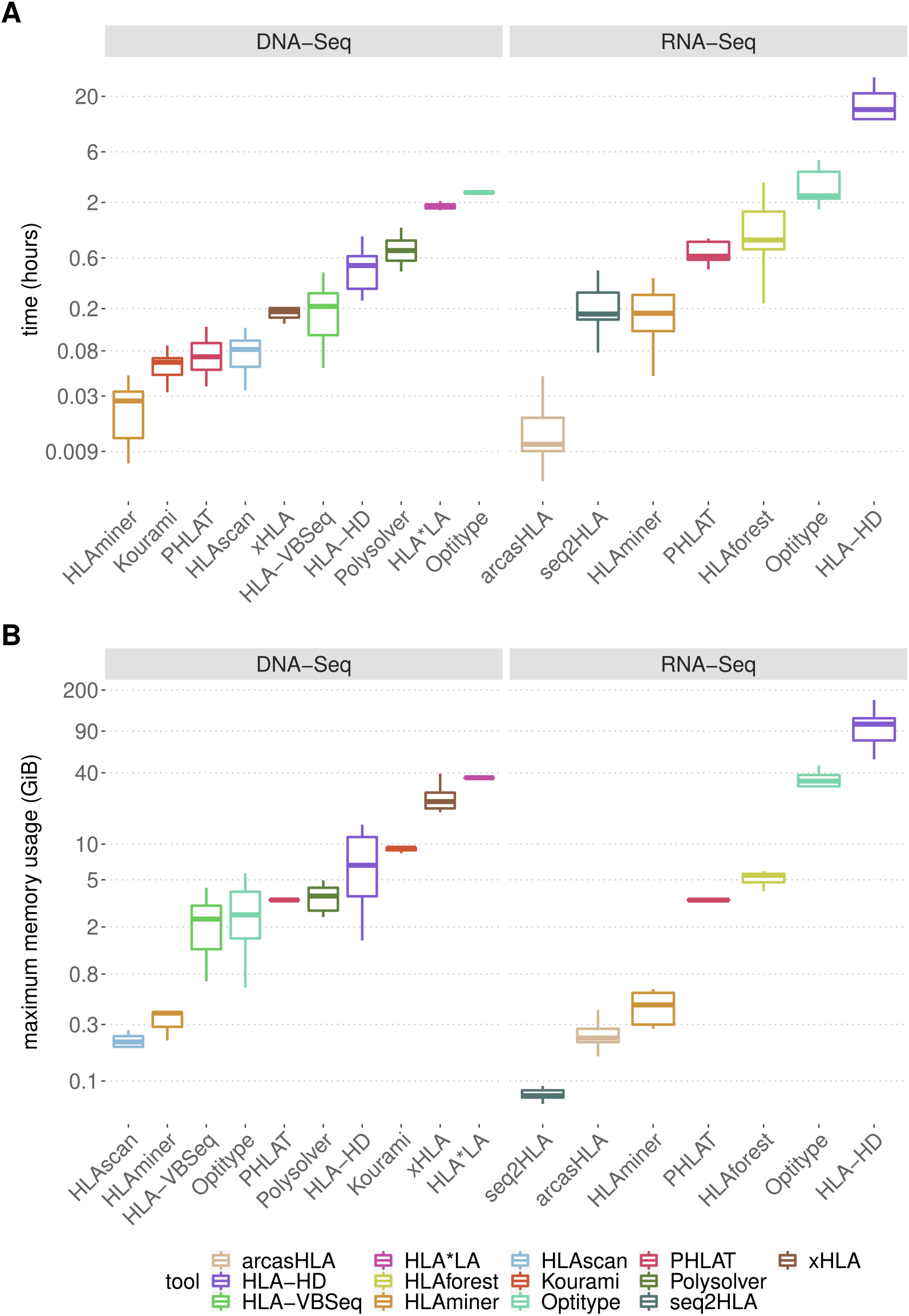
Computational resource consumption of the 13 selected tools. (**A-B**) Boxplots compare the resources needed by the different tools to analyse one sequencing file on a system with a single CPU core. Each tool was applied on DNA and/or RNA sequencing files *(n=10)*, as indicated at the top of the figure. Different tools are represented with a different colour of the boxplot, as indicated in the legend on the right. (**A**) Time consumption per sample. (**B**) Maximal memory consumption per sample.

Among the 10 DNA-supporting methods *Optitype* (median 2.48 hours) and *HLA*LA* (median 1.84 hours) require the largest computing time. The remaining DNA tools take less than 1 hour per file, with *HLAminer, Kourami* and *PHLAT* being the fastest (97s, 225s and 253s respectively). Apart from being computationally intensive, *HLA*LA* is also the most memory demanding DNA tool (median 36.3 GiB per file). Other DNA tools with a median memory consumption higher than 5 GiB are *xHLA* (median 22.9 GiB), *Kourami* (median 9.3 GiB) and *HLA-HD* (median 6.7 GiB). The relatively low memory usage of *Polysolver* makes it feasible to compensate for its long running time by processing multiple samples in parallel.

Among the 7 RNA-supporting methods, *HLA-HD* has the longest computing time per sample (median 15.0 hours). At the other end of the spectrum, the sole pseudoalignment-based tool *arcasHLA* takes only 38s per file. The most memory intensive tool is *HLA-HD* (median memory peaks of 103.1 GiB), followed by *Optitype* (median 34.1 GiB). The other RNA tools have a memory usage lower than 10 GiB. Remarkably, *HLAminer, PHLAT* and *HLA-HD*, which are compatible with both DNA and RNA data take a longer time on RNA data (median computing time per sample is 29.4, 8.9, 6.8 times longer for *HLA-HD, PHLAT* and *HLAminer* respectively).

### HLA*LA and HLA-HD are the best performing MHC class II genotyping tools on DNA data

The 10 selected algorithms that are compatible with DNA sequencing data were benchmarked using WES data from the *1000 Genomes project* [44] (Fig 2). Predictions were made for *HLA-A* (n = 1012), *HLA-B* (n = 1011), *HLA-C* (n = 1010), *HLA-DQB1* (n = 1008), *HLA-DRB1* (n = 1000) and *HLA-DQA1* (n = 68). *HLA-DPA1* and *HLA-DPB1* were not benchmarked due to the lack of available gold standard calls. For MHC-I genes (*HLA-A, HLA-B, HLA-C*), the best accuracy was obtained with *Optitype* (98.0%), followed by *Polysolver* and *HLA*LA* (94.9% and 94.4% respectively). For MHC-II genes (*HLA-DQA1, HLA-DQB1* and *HLA-DRB1*), the best allele predictions were made using *HLA-HD* and *HLA*LA* (96.2% and 95.7% accuracy respectively). These were the only two methods to reach an accuracy of 90% on all tested MHC-II genes. *HLAscan* (74.2%), *HLA-VBSeq* (60.2%) and *HLAminer* (53.8%) performed considerably worse than the other tools.

**Figure 2.**
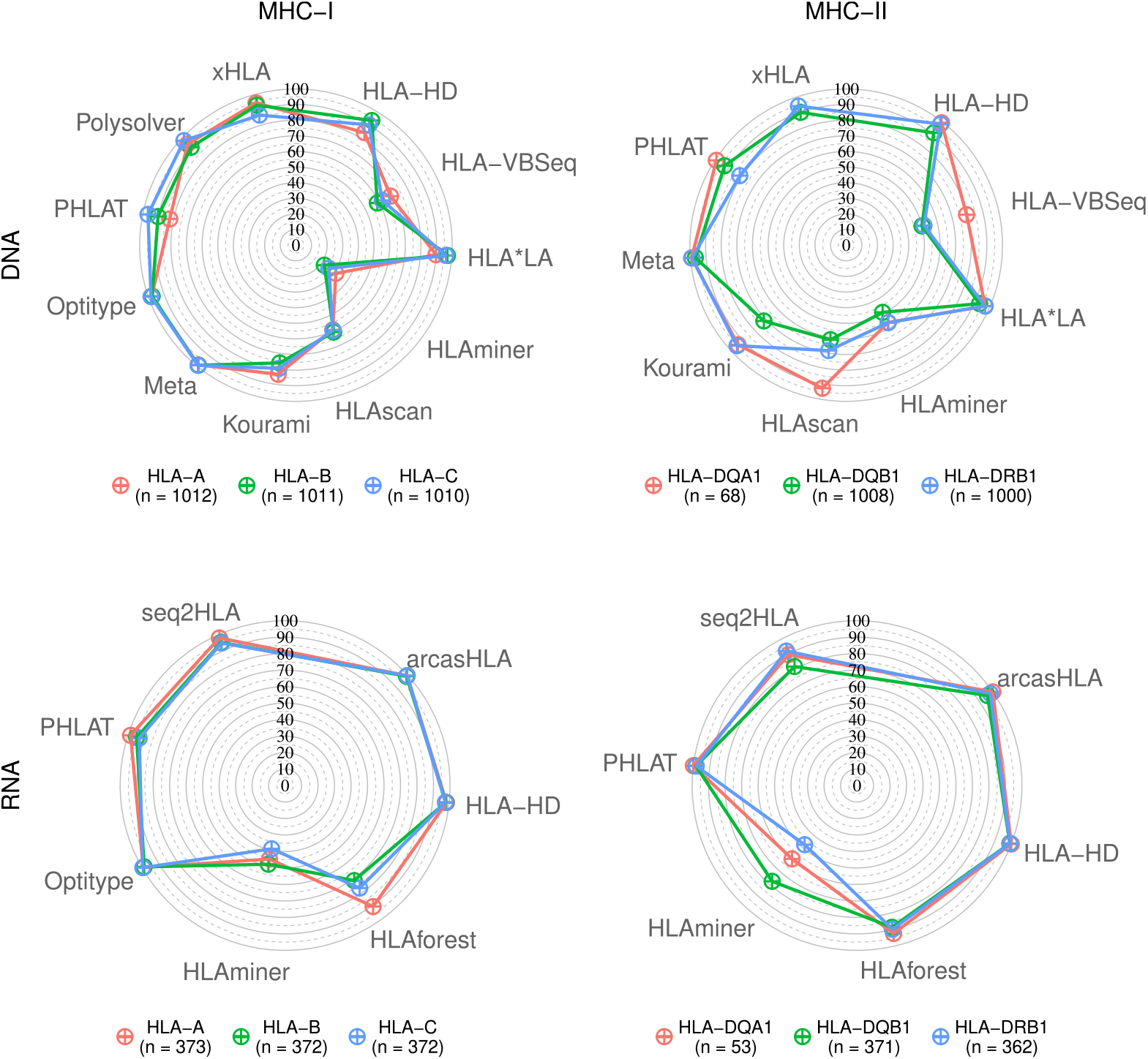
HLA allele prediction accuracies. Radar plots of HLA allele prediction accuracies on samples from the 1000 Genomes Project. Coloured lines represent different genes, as indicated in the legend below the plots. Corners of the radar plots correspond to the tools that were evaluated for that data type. The Meta tool correspond to the 4-tool consensus metaclassifier.

We observed large variabilities in calling accuracies between MHC class II genes (Fig 2). Overall, *HLA-DQB1* was the hardest MHC-II gene to call. Except for *PHLAT*, all tools obtained their worst MHC-II call accuracy on this gene. *HLA-DQA1*, on the other hand, was the gene with the highest calling accuracy for all tools that support it, except for *HLAminer* and *Kourami*.

Incorrect calls are either caused by wrong allele calls or a failure to make an allele call. When *HLAscan* and *Kourami* were able to make a call, their predictions were most often reliable (S1 Fig), but these tools regularly produced no output at all (S2 Fig). *HLA-VBSeq* and *HLAminer* had both a high rate of incorrect and failed calls (S1-S2 Figs).

Subsequently, we performed an independent benchmark using the smaller NCI-60 cell line dataset (n=58), which largely confirmed our results (S3 Fig). Additionally, this analysis indicated that the best performing MHC class II supporting tools also performed well on *HLA-DPB1*.

### HLA-HD, PHLAT and arcasHLA are the best performing MHC class II genotyping tools on RNA data

We then evaluated the 7 selected methods that support HLA calling on RNA sequencing data from the *1000 genomes project* [45] (Fig 2). Predictions were made for *HLA-A* (n = 373), *HLA-B* (n = 372), *HLA-C* (n = 372), *HLA-DQB1* (n = 371), *HLA-DRB1* (n = 362) and *HLA-DQA1* (n = 53).

A*rcasHLA* and *Optitype* had the best MHC-I allele predictions (99.4% and 99.2% accuracy, respectively), followed by *HLA-HD* (98.0%), *seq2HLA* (95.9%) and *PHLAT* (95.4%). Similar accuracies were found for MHC-II allele predictions, with *HLA-HD, PHLAT* and *arcasHLA* performing the best (99.4%, 98.9% and 98.1%, respectively). Contrary to its good prediction of MHC class I alleles, *seq2HLA* has a lower accuracy for MHC class II (87.8%).

The high MHC-I accuracies of *arcasHLA* and *Optitype* were confirmed on the independent NCI-60 dataset (91.8% and 90.0%, respectively; n=58). The accuracy of *HLA-HD, PHLAT* and *seq2HLA* was worse on the cell lines than in the benchmark on the 1000 genomes data (86.6%, 83.3% and 82.3%, respectively). As MHC-II is generally not expressed in cell lines, this benchmark was not performed for those genes.

### Correlation and concordance analyses on large independent datasets confirm the benchmarking results

Being one of the few large sequencing datasets for which gold standard HLA genotypes for both MHC classes are available, many algorithms included in our benchmark were developed, optimized and validated using files from the *1000 genomes* project, introducing a potential bias. Additionally, no evaluation was possible for *HLA-DPA1* and *HLA-DPB1*, due to the lack of gold standard HLA calls. Therefore, we performed an indirect and independent evaluation on a large NGS dataset obtained from TCGA (n=9162 and n=9761 for DNA and RNA respectively).

We first compared the observed allele frequencies for each tool with the expected population frequencies. We calculated how often each of the alleles was predicted by a certain tool to obtain an observed allele frequency, stratifying for Caucasian American (n= 7935) and African American (n=938) ethnicities. By comparing these frequencies to the expected allele frequencies, as derived from *Allele Frequency Net* [46], strong correlations (i.e., Pearson’s *r* 0.94 or higher for all genes and both populations) were found for the DNA-based tools *HLA-HD, HLA*LA, Optitype, Polysolver* and *xHLA* and for the RNA-based tools *Optitype, arcasHLA* and *PHLAT*. The correlations were considerably worse for *HLA-VBSeq, HLAminer* and *HLAforest* than for the other tools (Fig 3). These findings largely confirm the results of the benchmark on the 1000 genomes data. Notably, among the well performing tools, *arcasHLA* had a worse correlation for *HLA-DRB1*, which is mainly due to the discrepancy between the observed and predicted frequency of *HLA-DRB1*14:02* in this population (S4 Fig).

**Figure 3.**
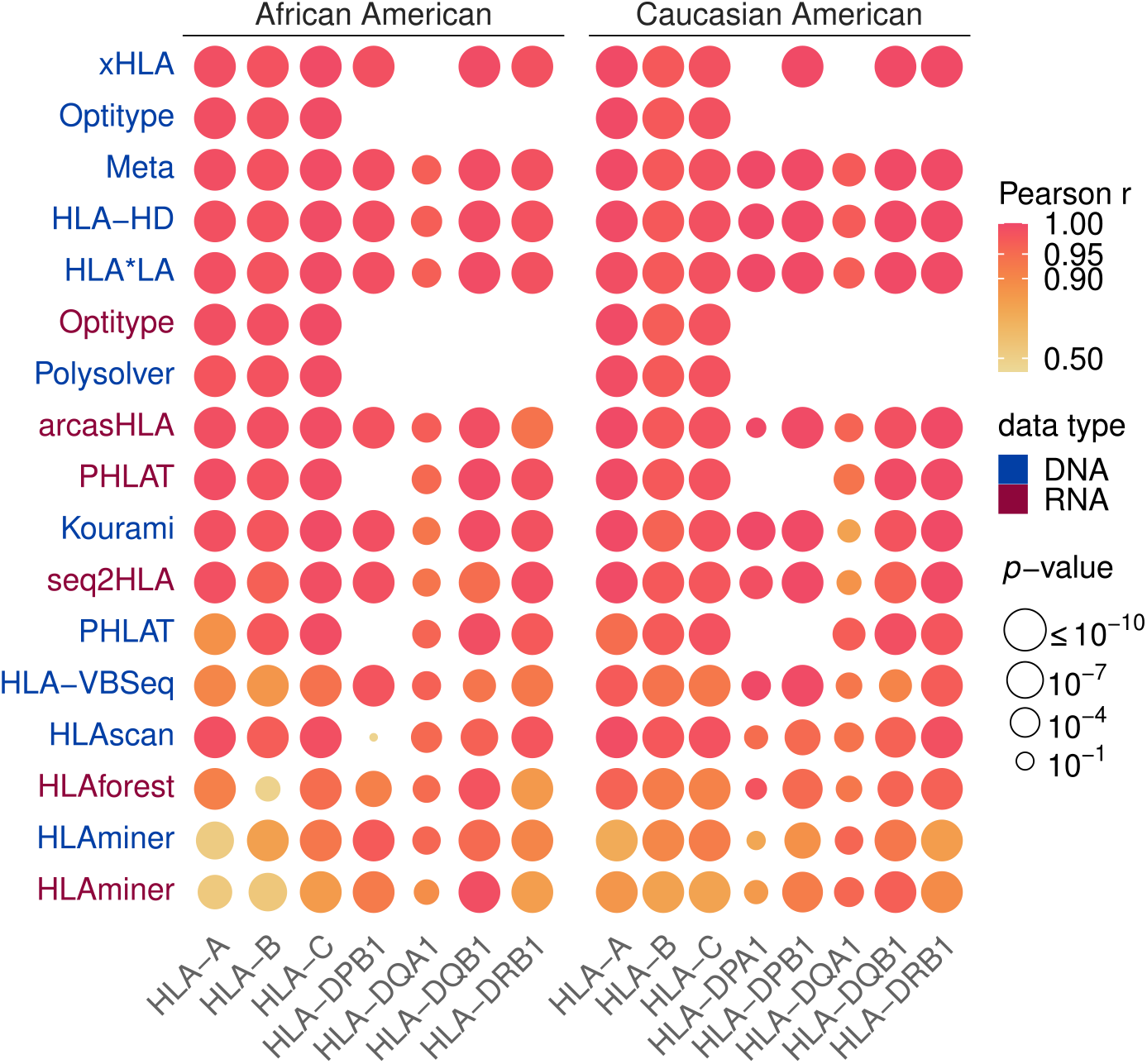
Correlations between observed and expected allele frequencies. Heatmap of correlations between observed allele frequencies and frequencies expected in an African American and in a Caucasian American population. Vertical axis indicates the tools, with different colours representing the data type (DNA or RNA) on which the tool was applied. Rows were sorted according to the mean correlation of the tool. Size of the circles indicates the *p*-value of the correlation test as indicated in legend. Absent circles indicate that the tool could not be evaluated on that gene.

We then calculated for each pair of tools how often their predictions are concordant (S5-S8 Figs). Tools that performed poorly in the previous analyses (e.g., *HLAminer, HLA-VBSeq* and *HLAforest*) consistently have a low concordance with all other tools. In contrary, tools that scored high in the previous analyses (such as *Optitype, HLA*LA, arcasHLA* and *HLA-HD*) made predictions that are consistent with each other. Noteworthy, this is also the case for *HLA-DPA1* and *HLA-DPB1*, two genes for which no gold standard data was available, suggesting that predictions for these genes are reliable as well.

### A consensus metaclassifier improves HLA predictions for DNA data

We noted that only for a very small fraction of the samples the genotypes are wrongly typed by all tools simultaneously (median 0.79% for DNA and 0.68% for RNA; Figures S9-S10). This complementarity of the tools’ allele predictions opens the possibility to combine predictions of different HLA callers into a consensus prediction. We first applied a majority voting algorithm to the output of all tools, with the predicted allele pair being the one with most votes. On the DNA data, this approach outperforms the predictions of each individual tool for all genes. This is best illustrated by the *HLA-DQB1* gene, where the accuracies increased from 93.2% with the best performing tool (*HLA*LA*) to 96.3% when the voting metaclassifier was used. On RNA data, where the best tools already attain accuracies over 99% by themselves, only minor improvements were made by combining the results (S11 Fig).

Based on these results, we determined the minimal number of tools that must be included in the DNA-based metaclassifier to produce reliable results (Fig 4). For the DNA data, including 4 tools in the model led to a considerable improvement for all genes for both MHC classes. The best accuracies were observed when *Optitype, HLA*LA, Kourami* and *Polysolver* were combined for MHC-I predictions (99.0% accuracy) and with *HLA*LA, HLA-HD, PHLAT* and *xHLA* for MHC-II predictions (98.4% accuracy). Raising the number of tools further only resulted in marginal gains. Strikingly, the accuracy of the *HLA-DQB1* allele predictions even decreases when more tools were included in the model. Therefore, we suggest combining the output of 4 tools for both MHC classes.

**Figure 4.**
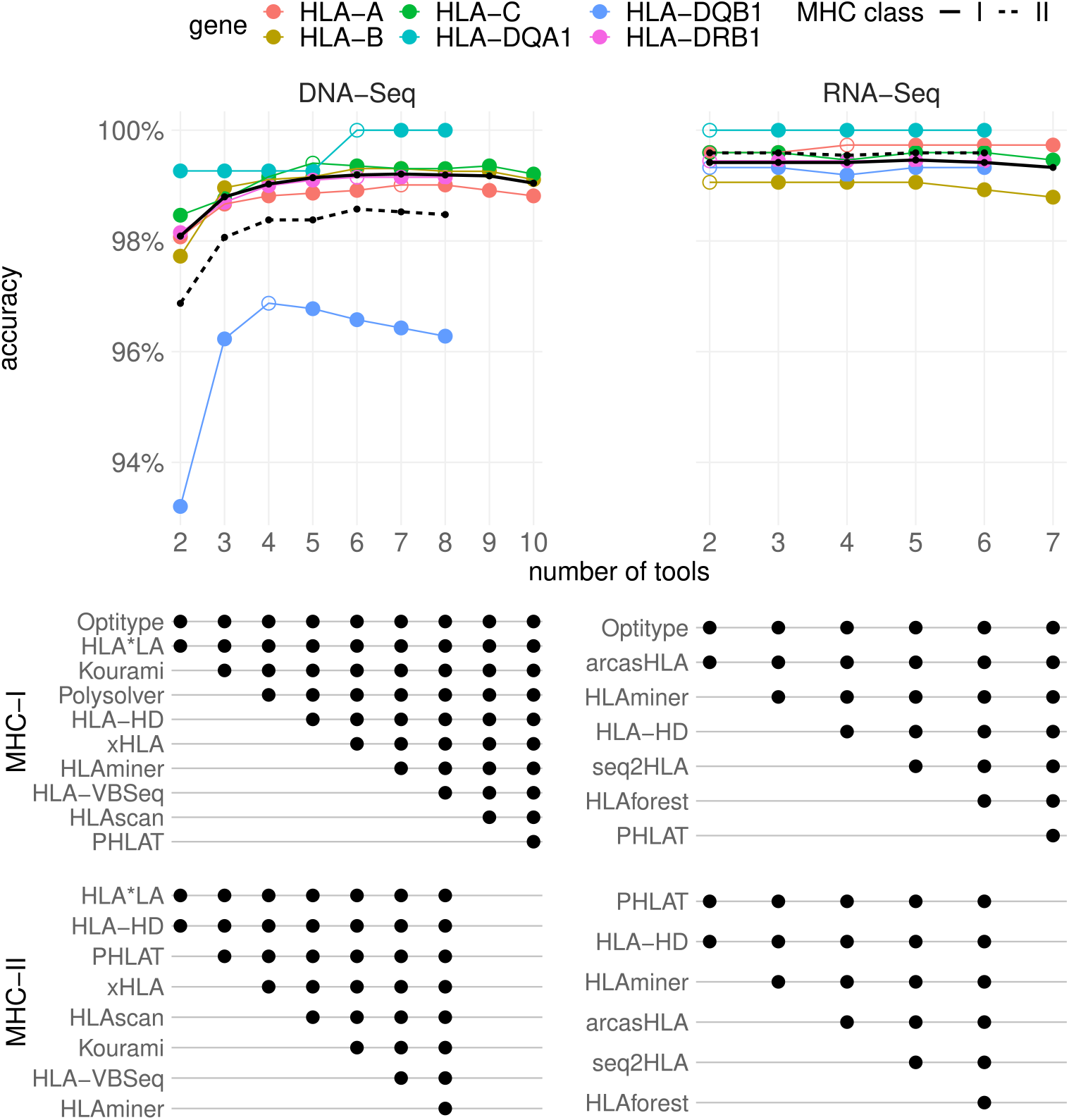
Accuracies of meta-prediction models with an increasing number of included tools. Tools were added one by one to the consensus metaclassifier model. At each step, the prediction accuracies of the best performing metaclassifier model for a given number of tools were plotted at the top of the figure. Unfilled markers are placed at the smallest number of tools where the maximal accuracy was obtained for that gene. Black lines indicate the average accuracy of the consensus predictions for the two MHC classes (averaged over all genes of that class). The table below the plot indicates which tools were selected in each model for a given number of tools.

To evaluate whether the good performance of this approach is generalizable to other datasets, we assessed the correlation between the expected allele frequencies and the allele frequencies observed using the 4-tool DNA consensus predictions on the TCGA dataset and compared the results with our previous findings. The allele frequencies predicted by the metaclassifier correlated better with the expected allele frequencies (Fig 3) than was the case for the individual tools that supported all genes of interest.

## Discussion

Rapid technological advancements in NGS have resulted in the generation of numerous publicly available WES and RNA-Seq datasets. These data have been critical for understanding the genomic basis of human carcinogenesis [47]. In the field of immuno-oncology, genomic data have also been used to study immune selection [48,49] and, additionally, the availability of corresponding clinical data opens possibilities for studying HLA-dependent cancer susceptibility or even differences in clinical ICB responses between cancer patients [12,50–52]. However, this requires that the HLA genotype for each subject can be accurately determined. An ever-increasing number of NGS-based HLA typing software applications have been developed. In this study, we benchmarked the performance of 13 publicly available tools. To our knowledge, this is the most extensive benchmark of MHC genotyping tools that has been performed so far (Table S4).

First, we evaluated the tools by comparing their output to genotypes derived from a PCR-based approach. While PCR methods are the gold standard for HLA typing, they have limitations that could lead to ambiguous typing results [53]. Furthermore, inconsistencies have been reported across PCR-based HLA typing datasets that are available for the 1000 genomes samples [54] which could have affected our benchmarking results. Therefore, we also used 2 other, indirect approaches to assess the performance of the different tools.

Both a concordance analysis between the tools’ predictions and a correlation analysis between predicted and expected allele frequencies confirmed our benchmarking results. To avoid biasing the results of this correlation analysis, we disabled ethnicity-specific allele frequencies for the algorithms that support this (i.e., *arcasHLA* and *Polysolver*). However, in the case of *arcasHLA*, when no specific ethnicity is specified, it uses prior frequencies that depend on the prevalence of the alleles in the entire human population, possibly hindering its ability to call alleles that are uncommon in the specified population. This is illustrated by the worse correlation between observed and expected allele frequencies of *arcasHLA* for *HLA-DRB1* in the African American population, due to an overestimation of the frequency of the rare *HLA-DRB1*14:02* allele.

We found that *Optitype, Polysolver, HLA-HD, HLA*LA* and *xHLA* are all solid choices for DNA-based MHC genotyping, while *Optitype, HLA-HD, arcasHLA* and *PHLAT* are the better performing tools for RNA data. On the other hand, *HLAminer, HLA-VBSeq* and *HLAScan* performed rather poorly in our benchmark. Similar trends were observed in previous independent benchmarking studies [14,17,19– 23] that focused on a subset of tools and/or genes (Table S4), with the exception of *xHLA* where we obtained considerably higher accuracies on WES data than reported in a study by Chen et al. [19]. The optimal strategy for HLA genotyping depends on a few factors: the availability of DNA or RNA data, the size of the dataset that needs to be analysed and the available computational resources. Additionally, MHC class II typing based on RNA data is only feasible on sequencing data derived from MHC-II expressing cells. For DNA data, *Optitype* and *HLA-HD* are the best performing individual tools for MHC class I and MHC class II typing, respectively. For RNA data, the same tools are recommended when sufficient computational resources are available. However, the large resource and time consumption of *HLA-HD* on RNA data makes its usage rather impractical on large datasets. As an alternative, *arcasHLA* is recommended, which is both the fastest and more accurate tool for RNA that supports all 5 MHC class II genes. Finally, we have demonstrated that the accuracy of the DNA-based HLA genotype predictions can be improved further by combining the output of *Optitype, HLA*LA, Kourami* and *Polysolver* for MHC-I typing and combining *HLA*LA, HLA-HD, PHLAT* and *xHLA* for MHC-II typing using a majority voting rule. For RNA data a similar approach did not lead to a further improvement of the prediction accuracies.

## Materials and Methods

### Selection of tools

A list of existing HLA genotyping tools for NGS data was compiled from literature between October and December 2020. The tools that fulfilled the following criteria were selected for further analysis: the tool should be free for academic use, support DNA and/or RNA sequencing data, should not require enrichment of the HLA region before sequencing and should be a Linux command line tool that we could successfully run on our system. When the authors provided instructions on how to update the IPD-IMGT/HLA database used by their tool, this database was updated to version 3.43. This was the case for three tools: *HLA-HD, HLAminer* and *Kourami*.

### Next-generation sequencing datasets for benchmark

Slices of the 1012 CRAM files of WES data from the *1000 Genomes on GRCh38* dataset [44] that were used for the benchmark on DNA data were obtained from the *International Genome Sample Resource* using the *samtools view* command (version 1.12). The following contigs were included in the download: the MHC region on the primary assembly (chr6:28509970-33480727), all 525 contigs starting with *HLA-* and all unmapped reads. The sliced BAM files for the RNA benchmark were obtained from the *Geuvadis* [45] RNA-Seq dataset (part of the *1000 genomes* project) via *ArrayExpress* (accession number *E-GEUV-1*). All reads mapped to the MHC region and the unmapped reads were included in the download. Sequencing data from NCI-60 cell lines [24] were obtained from the *Sequence Read Archive* with accession numbers *SRP150855* (WES) [55] and *SRP133178* (RNA) [56]. The NCI-60 sequencing data was realigned according to the same alignment pipeline used by the *1000 Genomes on GRCh38* dataset [44]: reads were aligned to the complete GRCh38 reference genome, including ALT contigs and HLA sequences, using an alternative scaffold-aware version of BWA-MEM. As done in the same 1000 genomes alignment pipeline, PCR-introduced duplicates were marked using the *markduplicates* function in BioBamBam (version 2.0.182). Aligned sequences of Whole Exome Sequencing (WES) and RNA sequencing experiments from *The Cancer Genome Atlas (TCGA)* were downloaded in BAM format from the *Genomic Data Commons (GDC)* portal. All 9162 available BAM files of Blood Derived normal WES samples were selected. For RNA-Seq, all 9762 RNA-Seq samples that were derived from primary tumours and were aligned using the “STAR 2-Pass” workflow, were selected. Reads mapped to the MHC region of chromosome 6 (chr6:28509970-33480727) and unmapped reads were extracted from the BAM files and downloaded following the instructions that are described in the GDC API. For the RNA-Seq samples one file failed to download after multiple attempts. The resulting dataset consists of 9162 blood-derived normal WES samples and 9761 primary tumour RNA-Seq samples from 33 available cancer types. The most resource intensive RNA tools were applied on a subset of the TCGA dataset. *Optitype* was applied on 2226 RNA files, *HLAforest* on 2900 files and *HLA-HD* was not applied on the TCGA data.

### Gold standard HLA typing data

Gold standard PCR-based HLA calls for the samples from the *1000 genomes on GRCh38* dataset were provided by three earlier studies [26–28]. The HLA genotypes from these datasets were merged. Where the calls did not agree, the calls by Gourraud et al. [57] were preferred. For the NCI-60 cell lines, PCR-based HLA genotypes were provided in a study by Adams et al [29]. For both reference datasets alleles were mapped to the corresponding G-groups, as defined by IPD-IMGT (http://hla.alleles.org/alleles/g_groups.html), and trimmed to the second-field resolution.

### HLA allele predictions

All 13 selected tools were run on the sliced BAM files following the guidelines of the authors. For tools requiring FASTQ input files, a FASTQ file was extracted from the sliced BAM files using *samtools fastq*. For *HLAScan*, which supports input files in either file format, the input was provided in BAM format. For tools that allowed to specify a list of loci that should be called: *HLA-A, HLA-B, HLA-C, HLA-DPA1, HLA-DPB1, HLA-DQA1, HLA-DQB1* and *HLA-DRB1* were chosen. *Kourami* was run with the *-a* (additional loci) parameter to call the *HLA-DPA1* and *HLA-DPB1* genes. In rare cases, this led to a crash of the tool and *Kourami* was run again without the *-a* parameter. For *HLAminer* only the HPRA mode was evaluated. *xHLA, Polysolver* and *HLA-VBSeq* were not compatible with BAM files that are aligned to a reference genome build that includes alternative contigs. For these tools, an additional realignment step was performed before the tool was executed. Input data for *xHLA* and *Polysolver* were realigned to a GRCh38 build that excludes alternative contigs. The input data for *HLA-VBSeq* was realigned to GRCh37. All allele predictions were mapped to the corresponding G-groups and trimmed at second-field resolution.

### Measuring the resource consumption

The running time and memory consumption required by the tools were measured for a random subset of 10 DNA and 10 RNA sequencing files from the TCGA project. Each tool was executed in a separate Docker container (version 19.03.3) that was allocated a single CPU core. When the package provided a parameter to specify the number of threads, this was set to 1. Per file, the memory usage of the Docker container was monitored using the *docker stats* command. The running time was calculated as the time interval between the start and the end of the tool, excluding the time to start the Docker container. Pre-processing steps related to realignment to a different genome build (as required for *xHLA, Polysolver* and *HLA-VBSeq*) were not included in the resource consumption assessment. For HLA-HD the analysis of a single sample did not complete successfully as the required amount of memory exceeded what we have available on our system.

### Performance metric

For each sample, two allele predictions were made. An allele prediction was labelled “correct” when it was listed as one of the two alleles in the gold standard for that patient. When a tool made a homozygous prediction, while the gold standard was heterozygous, at most one of the two predictions was labelled “correct” for that sample. The accuracy of the predictions is then defined as the proportion of all correctly predicted alleles divided by twice the number of samples. Samples where the gold standard was missing for a particular gene were ignored for that gene.

### Population frequency data

Lists of expected HLA allele frequencies for an African American and for a Caucasian American population were constructed based on 18 different studies in the *Allele Frequency Net* [46] database (Table S3). The studies were selected based on the following criteria. First, we required that the study was conducted on a *Black* or *Caucasoid* population from the United States. This was not possible for *HLA-DPA1* where no HLA allele frequencies were available for these ethnicities. As a substitute, the allele frequencies of three European populations (French, Swedish and Basques) were used to approximate the allele frequencies for this gene in Caucasian Americans. As a second requirement, the HLA calls should be determined by a PCR-based method. Thirdly, the *Allele Frequency Net* database should have assigned a gold label (i.e., allele frequency sums to 1, sample size of study > 50, and at least 2-field resolution) to the study for the gene of interest. Lastly, it was required that the subjects included in the selected studies were healthy subjects (i.e., selected for an anthropological study, blood donors, bone marrow registry or controls for a disease study). Allele frequencies from different studies were combined by taking the average frequency, weighted according to the study’s sample size. All alleles were mapped to the corresponding G-groups and trimmed at second-field resolution.

### Correlation between expected and observed allele frequencies

For all tools and for each supported data type, the number of times that each allele was called was counted. This count was divided by the total number of samples to obtain the “observed allele frequency”. The Pearson correlation was calculated between observed allele frequencies and the allele frequencies that were expected based on the *Allele Frequency Net* database.

### Concordance of predictions among different tools

Per gene, the concordance of the predictions between each pair of tools was assessed by counting the number of allele pair predictions made by the first tool that were also made by the second tool (for the same sample and gene). Samples where one of both tools did not make a prediction were not considered. This analysis was performed on the 1000 genomes and TCGA dataset.

### Consensus HLA predictions

A majority voting rule was used to determine the most likely HLA genotype for each sample. For each gene of interest, we selected the pair of alleles that has been predicted the most frequently for that sample (i.e., outputted by the highest number of tools). When ties occurred (i.e., multiple allele pairs had equal numbers of predictions), priority was given to the allele pair that was predicted by the tool with the best individual performance for that gene.

### Selecting a minimum number of tools to make consensus HLA predictions

The minimal set of tools that must be included in the majority voting scheme to make reliable consensus predictions was determined using an iterative procedure. Initially, two tools were selected for the model: the tool that performed the best in the benchmark on the 1000 genomes data and the one that best complements that tool. The latter tool was defined as the tool that most often made a correct prediction (for both alleles) on the samples that were wrongly predicted by the best performing tool. Tools were added one by one to this initial model. At each step, the consensus predictions were made and evaluated using the gold standard HLA calls. The tool that led to the greatest increase in accuracy was added to the model. This procedure was repeated until all tools were selected.

### Hardware and software environment

Analyses were performed on Ubuntu 20.04 on a Dell EMC PowerEdge R940xa server with 4 Intel Xeon Gold 6240 CPUs (2.60 GHz), each with 18 physical CPU cores, and 376 GiB RAM installed.

### Data processing and statistical analysis

Data processing and statistical analyses were performed using R (version 4.0).

## Supporting information

**S1 Fig. Fraction of correct allele predictions (1000 genomes)**. Radar plots depicting the fraction of correct allele predictions relative to the total number of alleles for which the algorithm was able to make a prediction on the 1000 genomes dataset. Coloured lines represent different genes, as indicated in the legend below the plots. Corners of the radar plots correspond to the tools that were evaluated for that data type. The Meta tools correspond to the 4-tools metaclassifiers.

**S2 Fig. Fraction of successful allele predictions (1000 genomes)**. Radar plots depicting the fraction of alleles for which the tool was able to make a prediction on the 1000 genomes dataset. Coloured lines represent different genes, as indicated in the legend below the plots. Corners of the radar plots correspond to the tools that were evaluated for that data type. The Meta tools correspond to the 4-tools metaclassifiers.

**S3 Fig. HLA allele prediction accuracies on NCI-60 cell lines**. Radar plots of HLA allele prediction accuracies on data from NCI-60 cell lines. Coloured lines represent different genes, as indicated in the legend below the plots. Corners of the radar plots correspond to the tools that were evaluated for that data type.

**S4 Fig. Expected frequency of HLA-DRB1 alleles in an African American population vs frequencies predicted by arcasHLA**. Scatter plot that compares the allele frequency as predicted by arcasHLA (x-axis) with the expected allele frequencies based on data from *Allele Frequency Net* (y-axis).

**S5 Fig. Concordance of HLA calls between each pair of tools on DNA data (1000 genomes)**. Heatmaps representing the concordance of the HLA calls between each pair of tools, applied on the 1000 genomes DNA data. Hierarchical clustering was applied on the tools. The Meta tool corresponds to the 4-tool consensus metaclassifier.

**S6 Fig. Concordance of HLA calls between each pair of tools on RNA data (1000 genomes)**. Heatmaps representing the concordance of the HLA calls between each pair of tools, applied on the 1000 genomes RNA data. Hierarchical clustering was applied on the tools.

**S7 Fig. Concordance of HLA calls between each pair of tools on DNA data (TCGA)**. Heatmaps representing the concordance of the HLA calls between each pair of tools, applied on the TCGA DNA data. Hierarchical clustering was applied on the tools. The Meta tool corresponds to the 4-tool consensus metaclassifier.

**S8 Fig. Concordance of HLA calls between each pair of tools on RNA data (TCGA)**. Heatmaps representing the concordance of the HLA calls between each pair of tools, applied on the TCGA RNA data. Hierarchical clustering was applied on the tools.

**S9 Fig. Correctness of predictions on DNA data**. Heatmap indicating correctness of predictions on DNA data for each sample (rows) and tool (columns). Hierarchical clustering was applied on tools and samples. Dendrogram for the tools is shown on top of the plots. Dendrogram for the samples is shown right of the plots.

**S10 Fig. Correctness of predictions on RNA data**. Heatmap indicating correctness of predictions on RNA data for each sample (rows) and tool (columns). Hierarchical clustering was applied on tools and samples. Dendrogram for the tools is shown on top of the plots. Dendrogram for the samples is shown right of the plots.

**S11 Fig. Comparison of accuracies of all-tool metaclassifier with best performing individual tool per gene**. Barplots comparing the accuracy of the best tool for each gene and data type to the accuracy of a classifier that chooses an HLA genotype from the output of all tools that support that data type and gene based on a majority voting rule. Bars in a red correspond to the accuracies of the voting classifier. Bars in blue correspond to the accuracies of the best individual tool for that gene.

**S1 Table. Motivation for exclusion of 9 tools from the study**. Overview of tools that were not benchmarked in our study and the reason for their exclusion. Excluded tools were either not *freely available for academic use*, do not use *FASTQ or BAM input files from WGS, WES and/or RNA-Seq* experiments (e.g., enrichment of the HLA region prior to sequencing is needed) or we were not able to get them *running on Ubuntu 20*.*04*.

**S2 Table. Algorithmic characteristics of the 13 selected tools**. Table that compares the 13 selected HLA genotyping algorithms. Methods differ in the method to align reads to the HLA allele reference sequences (alignment method) and in the subsequent allele prioritization approach (allele pair prioritization). The column *tool-specific steps* describes additional particularities of the tools.

**S3 Table. Overview of AFN datasets used for expected allele frequencies**. Overview of populations (studies) from the *Allele Frequency Net database* which were used to compile the list of expected HLA allele frequencies.

**S4 Table. Comparison of our results with 7 other independent benchmark studies**. The allele prediction accuracies obtained in our benchmark on the 1000 genomes data are compared with 7 other independent benchmark papers. Each score was assigned a colour code: red under 50%, yellow between 50% and 80% and green above 80%. A grey box indicates that a tool was not evaluated for the corresponding gene in that study. The second row of the table indicates the gene or MHC class, where *class I + II* corresponds to the overall (combined) accuracy for MHC class I and class II. For the Lee, Kiyotani, Bauer and Yu studies the tools were evaluated on data from the 1000 genomes project. The studies by Liu, Yi and Chen used a different in-house dataset. DNA data always refers to WES data.

## Code and Data Availability

The code used to produce the results described in this manuscript are available at https://github.com/CCGGlab/mhc_genotyping.

## Acknowledgments

The results shown here are in whole or part based upon data generated by the TCGA Research Network: https://www.cancer.gov/tcga.

