## Supplementary figures S1-S11 for "Benchmark of tools for *in silico* prediction of MHC class I and class II genotypes from NGS data"

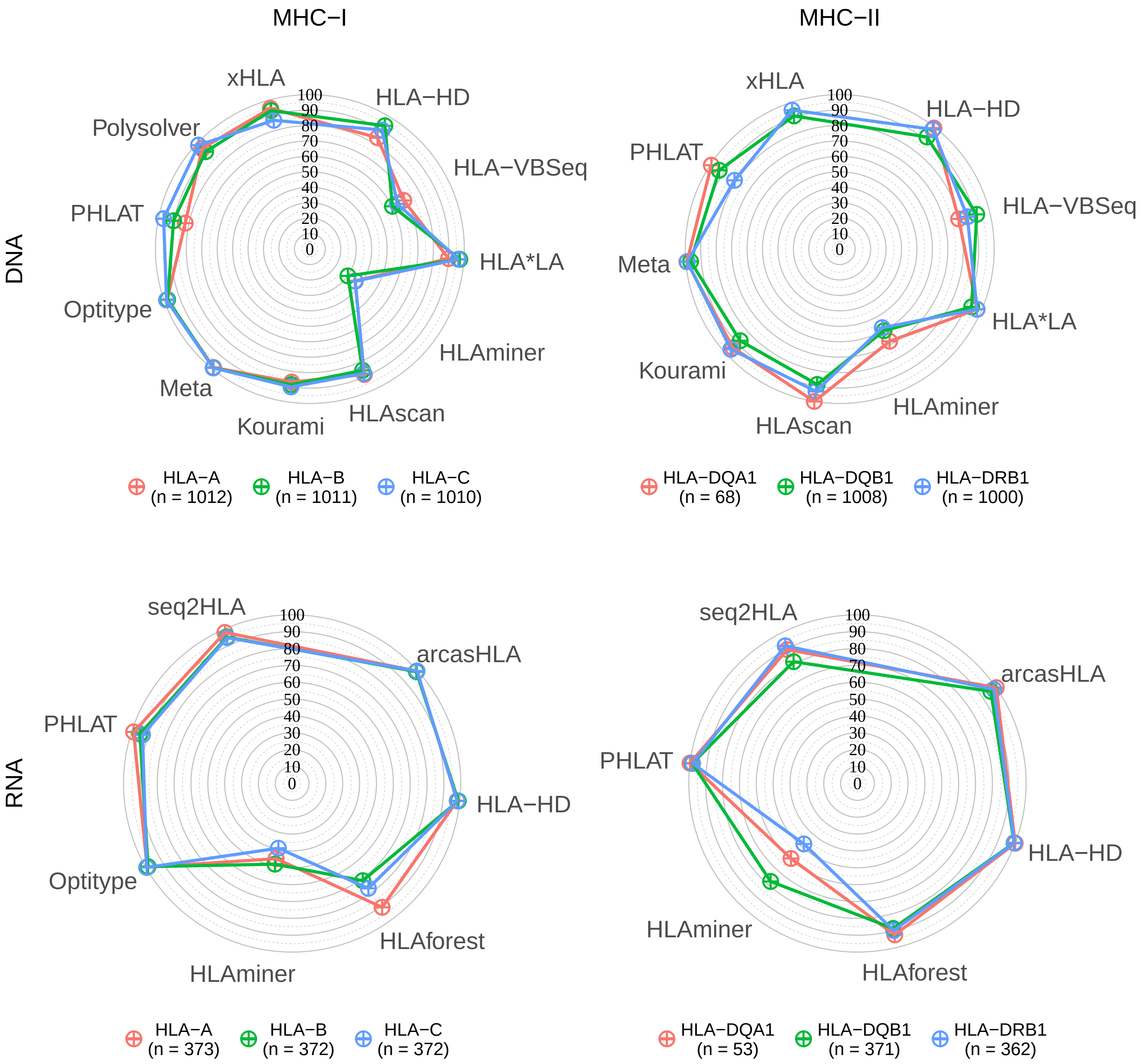


**Figure S1: Fraction of correct allele predictions (1000 genomes)**
Radar plots depicting the fraction of correct allele predictions relative to the total number of alleles for which the algorithm was able to make a prediction on the 1000 genomes dataset. Coloured lines represent different genes, as indicated in the legend below the plots. Corners of the radar plots correspond to the tools that were evaluated for that data type. The Meta tools correspond to the 4-tools metaclassifiers.

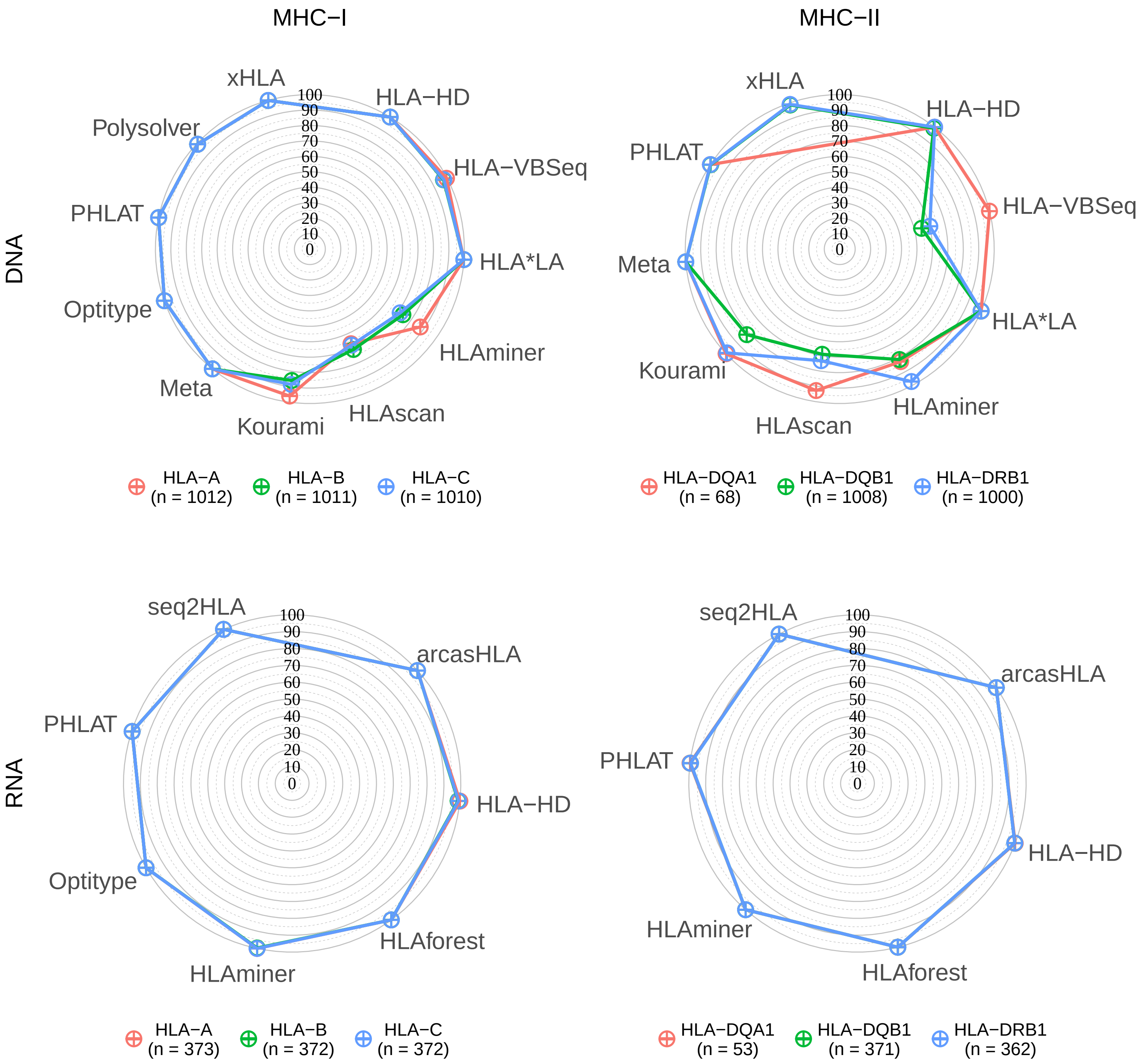


**Figure S2: Fraction of successful allele predictions (1000 genomes)**
Radar plots depicting the fraction of alleles for which the tool was able to make a prediction on the 1000 genomes dataset. Coloured lines represent different genes, as indicated in the legend below the plots. Corners of the radar plots correspond to the tools that were evaluated for that data type. The Meta tools correspond to the 4-tools metaclassifiers.

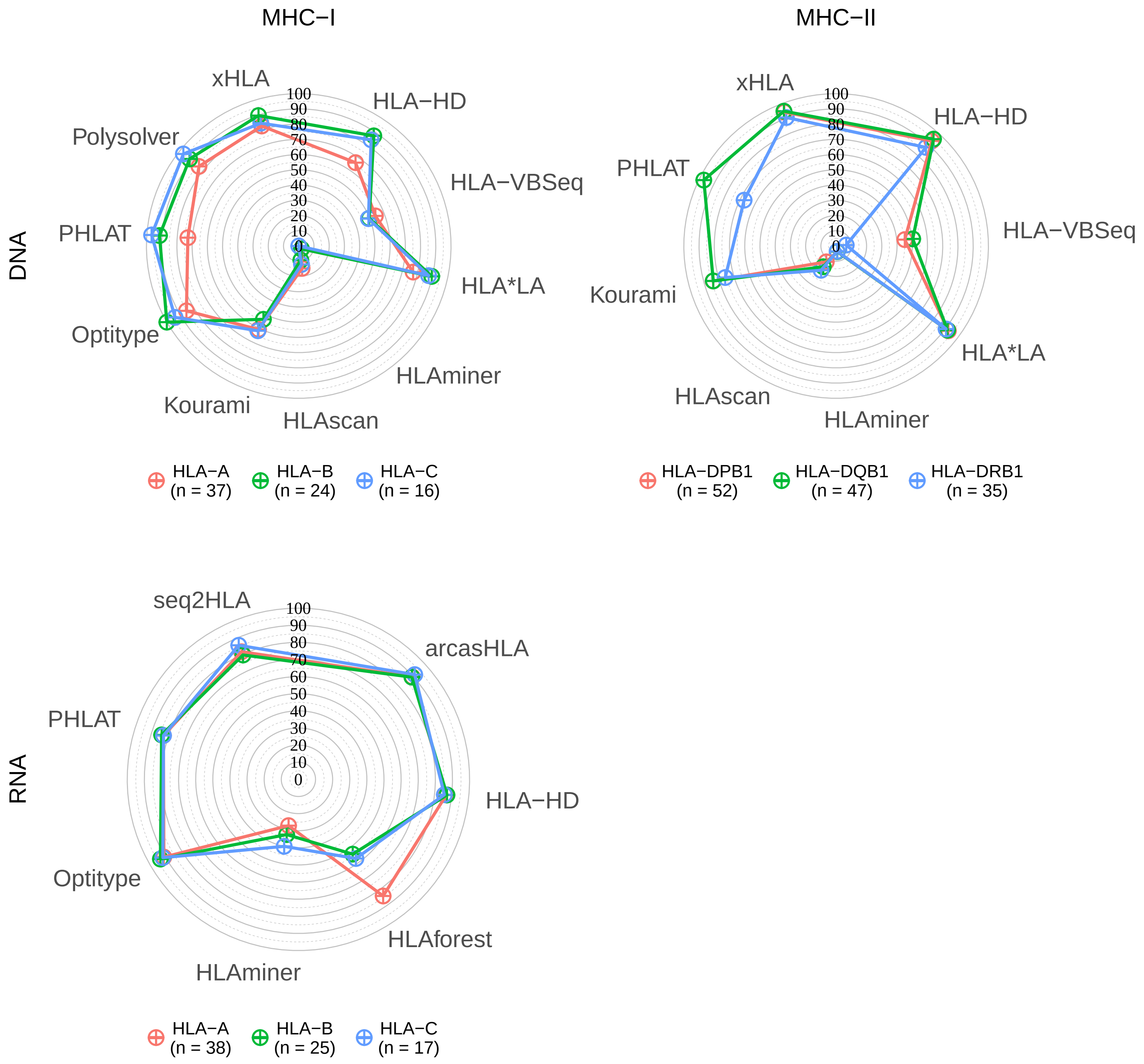


**Figure S3: HLA allele prediction accuracies on NCI-60 cell lines**
Radar plots of HLA allele prediction accuracies on data from NCI-60 cell lines. Coloured lines represent different genes, as indicated in the legend below the plots. Corners of the radar plots correspond to the tools that were evaluated for that data type.

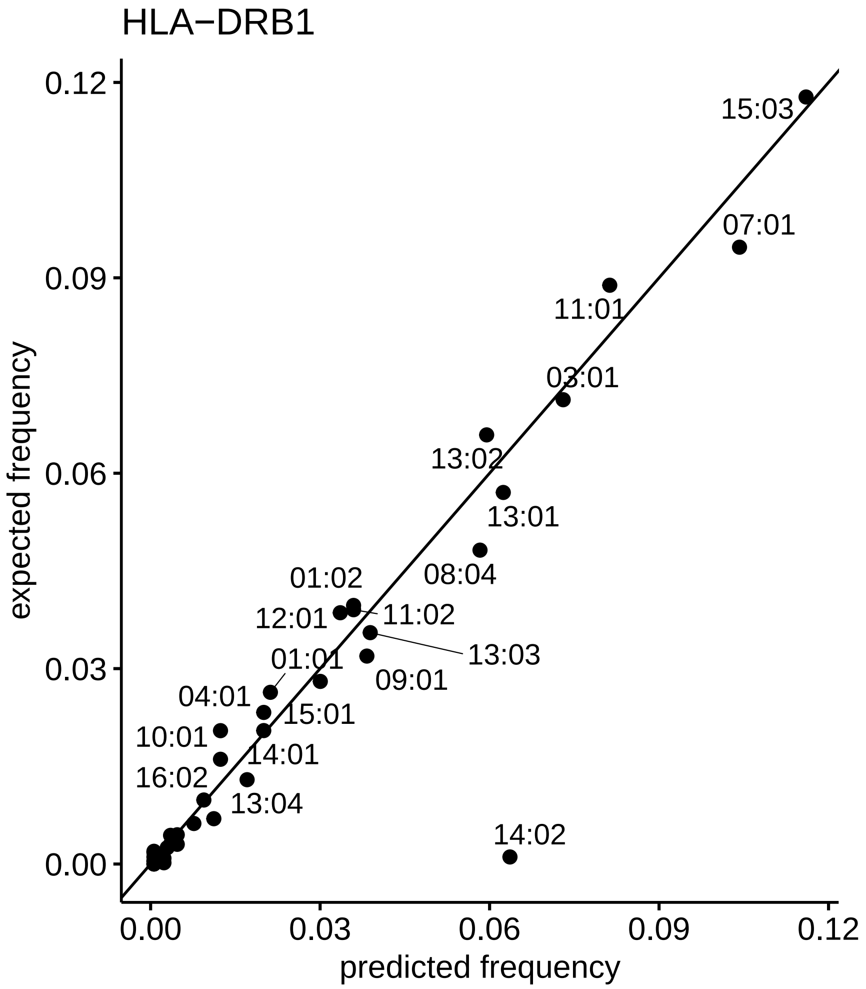


**Figure S4: Expected frequency of HLA-DRB1 alleles in an African American population vs frequencies predicted by arcasHLA**

Scatter plot that compares the allele frequency as predicted by arcasHLA (x-axis) with the expected allele frequencies based on data from Allele Frequency Net (y-axis).

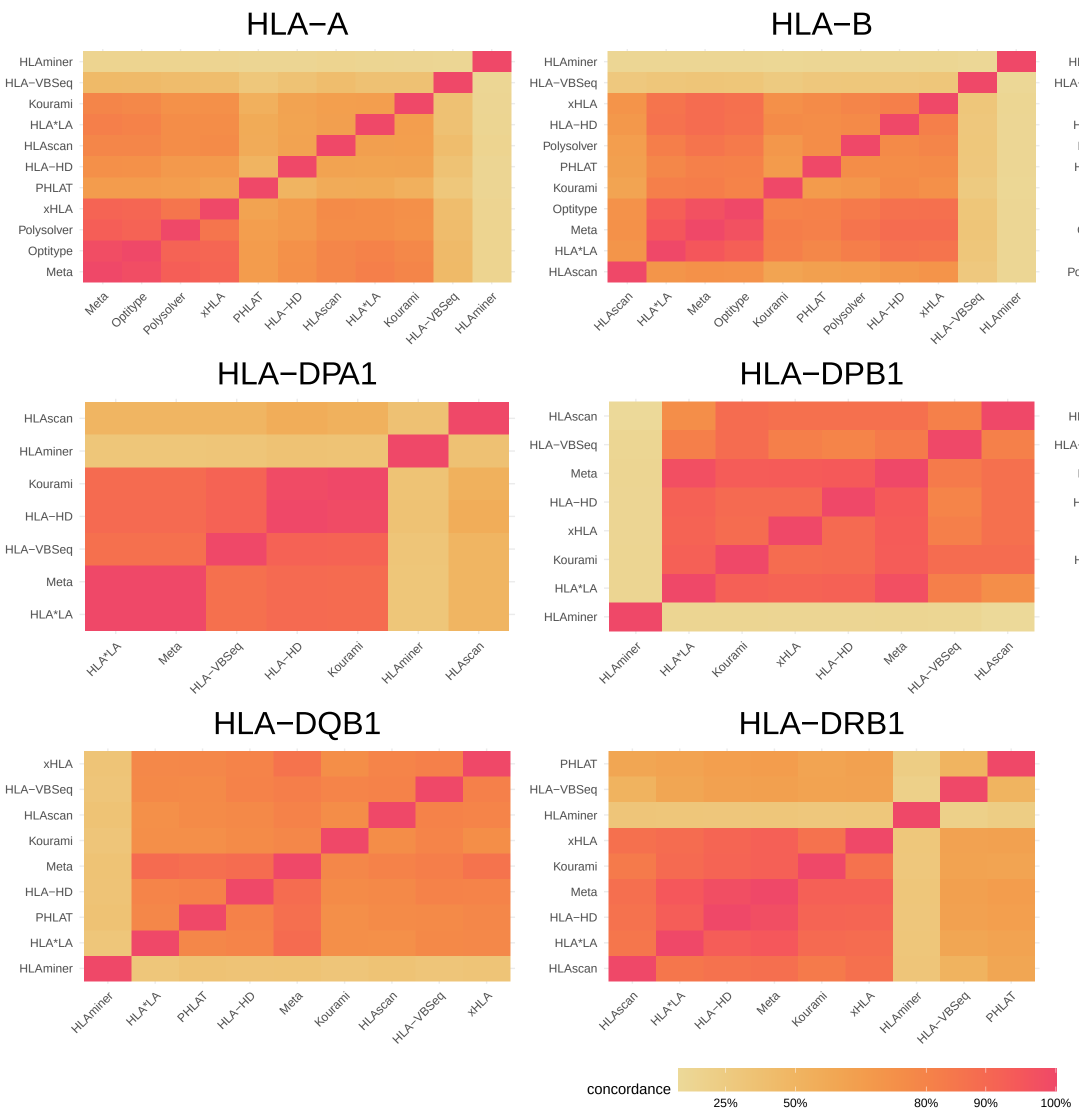


**Figure S5: Concordance of HLA calls between each pair of tools on DNA data (1000 genomes)** Heatmaps representing the concordance of the HLA calls between each pair of tools, applied on the 1000 genomes DNA data. Hierarchical clustering was applied on the tools. The Meta tool corresponds to the 4-tool consensus metaclassifier.

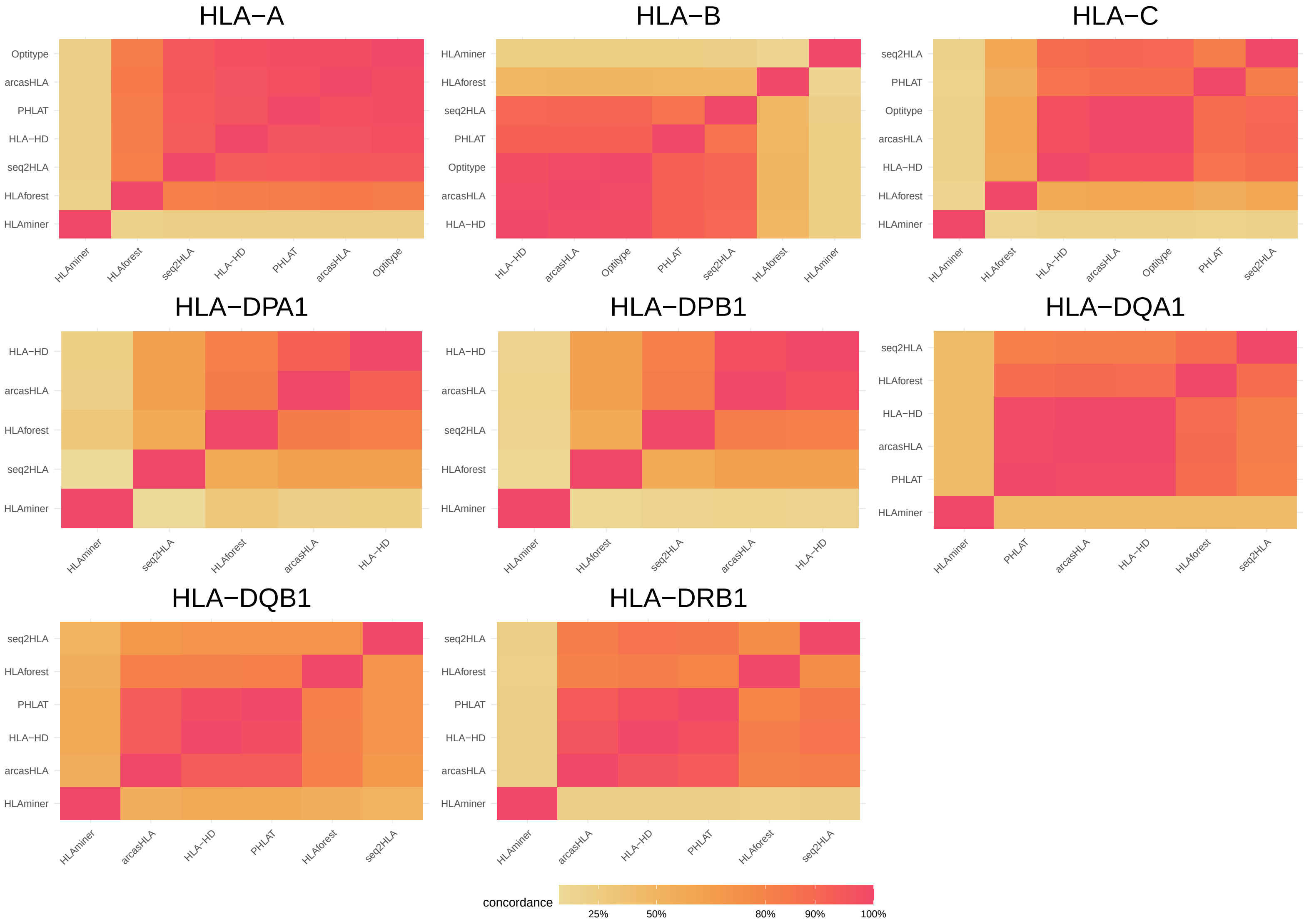


**Figure S6: Concordance of HLA calls between each pair of tools on RNA data (1000 genomes)** Heatmaps representing the concordance of the HLA calls between each pair of tools, applied on the 1000 genomes RNA data. Hierarchical clustering was applied on the tools.

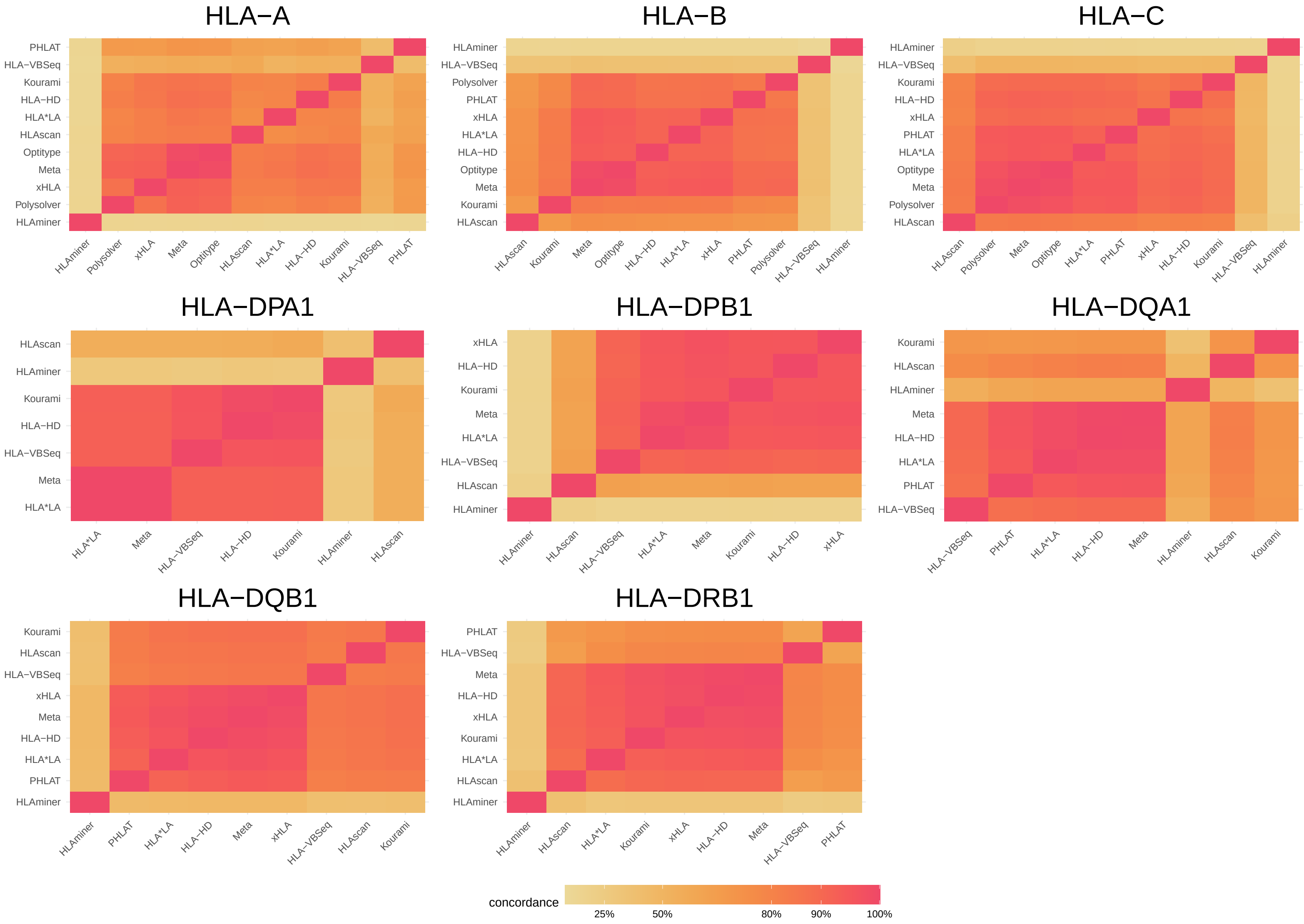


**Figure S7: Concordance of HLA calls between each pair of tools on DNA data (TCGA)** Heatmaps representing the concordance of the HLA calls between each pair of tools, applied on the TCGA DNA data. Hierarchical clustering was applied on the tools. The Meta tool corresponds to the 4-tool consensus metaclassifier.

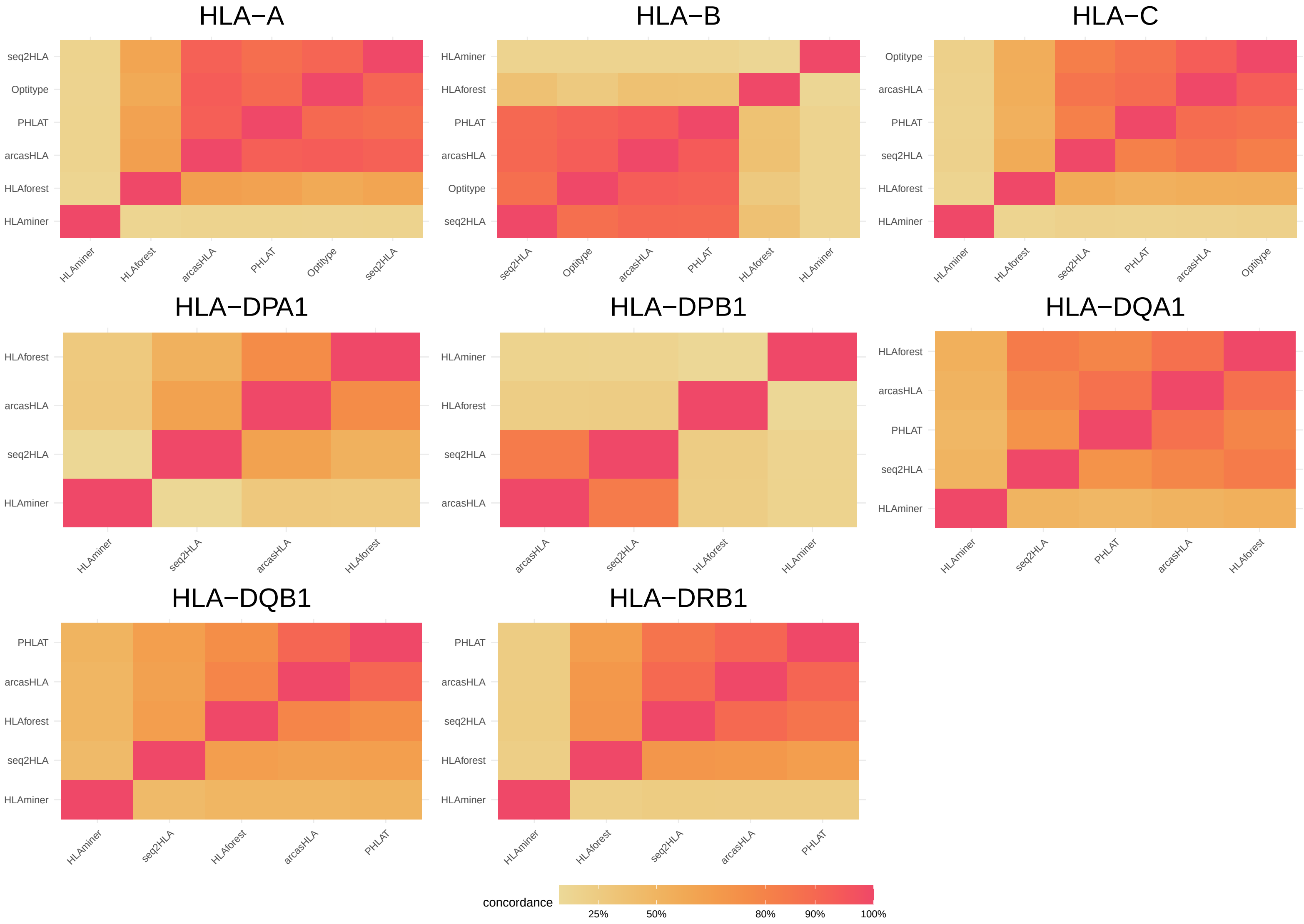


**Figure S8: Concordance of HLA calls between each pair of tools on RNA data (TCGA)**
Heatmaps representing the concordance of the HLA calls between each pair of tools, applied on the TCGA RNA data. Hierarchical clustering was applied on the tools.

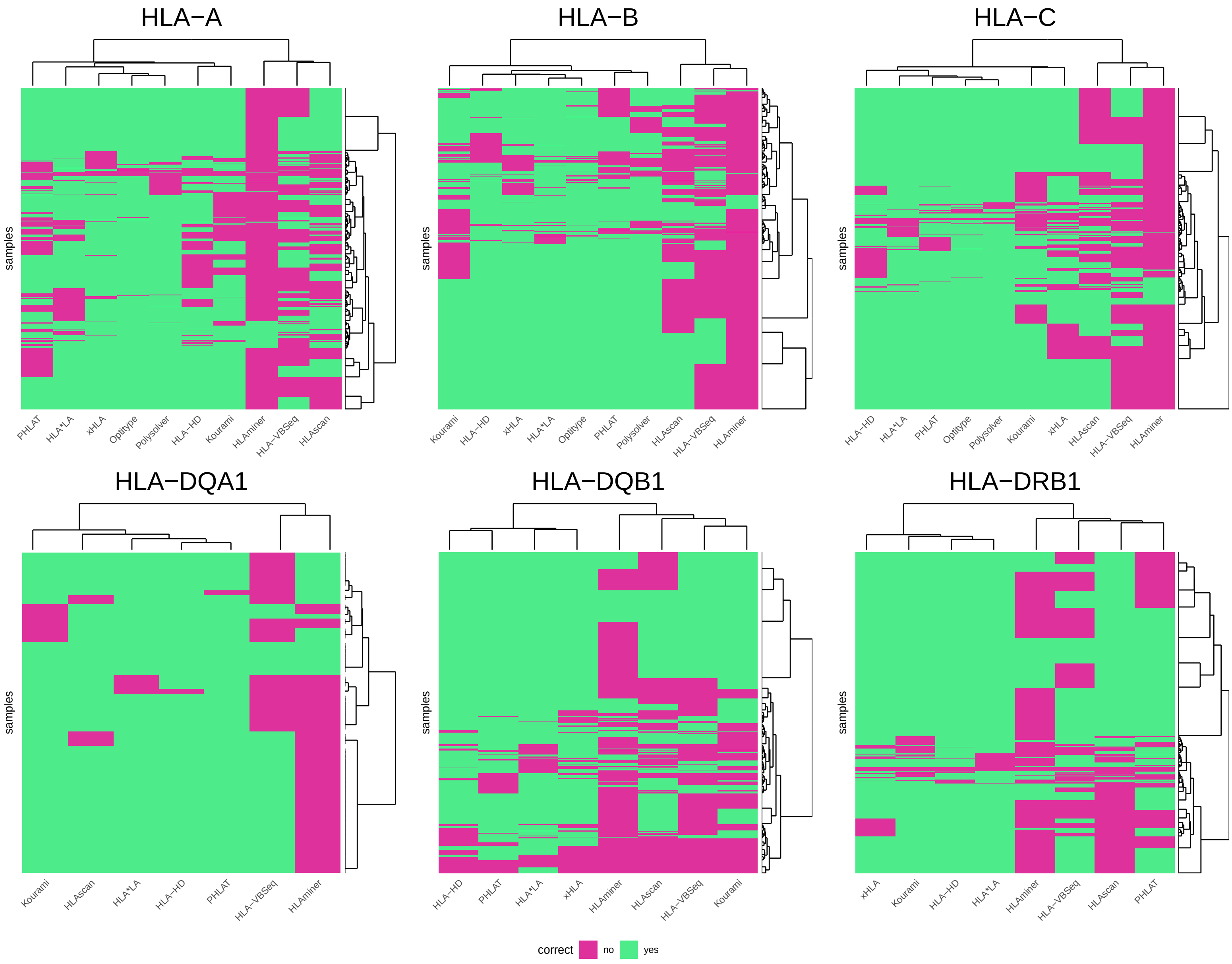


**Figure S9: Correctness of predictions on DNA data**
Heatmap indicating correctness of predictions on DNA data for each sample (rows) and tool (columns). Hierarchical clustering was applied on tools and samples. Dendrogram for the tools is shown on top of the plots. Dendrogram for the samples is shown right of the plots.

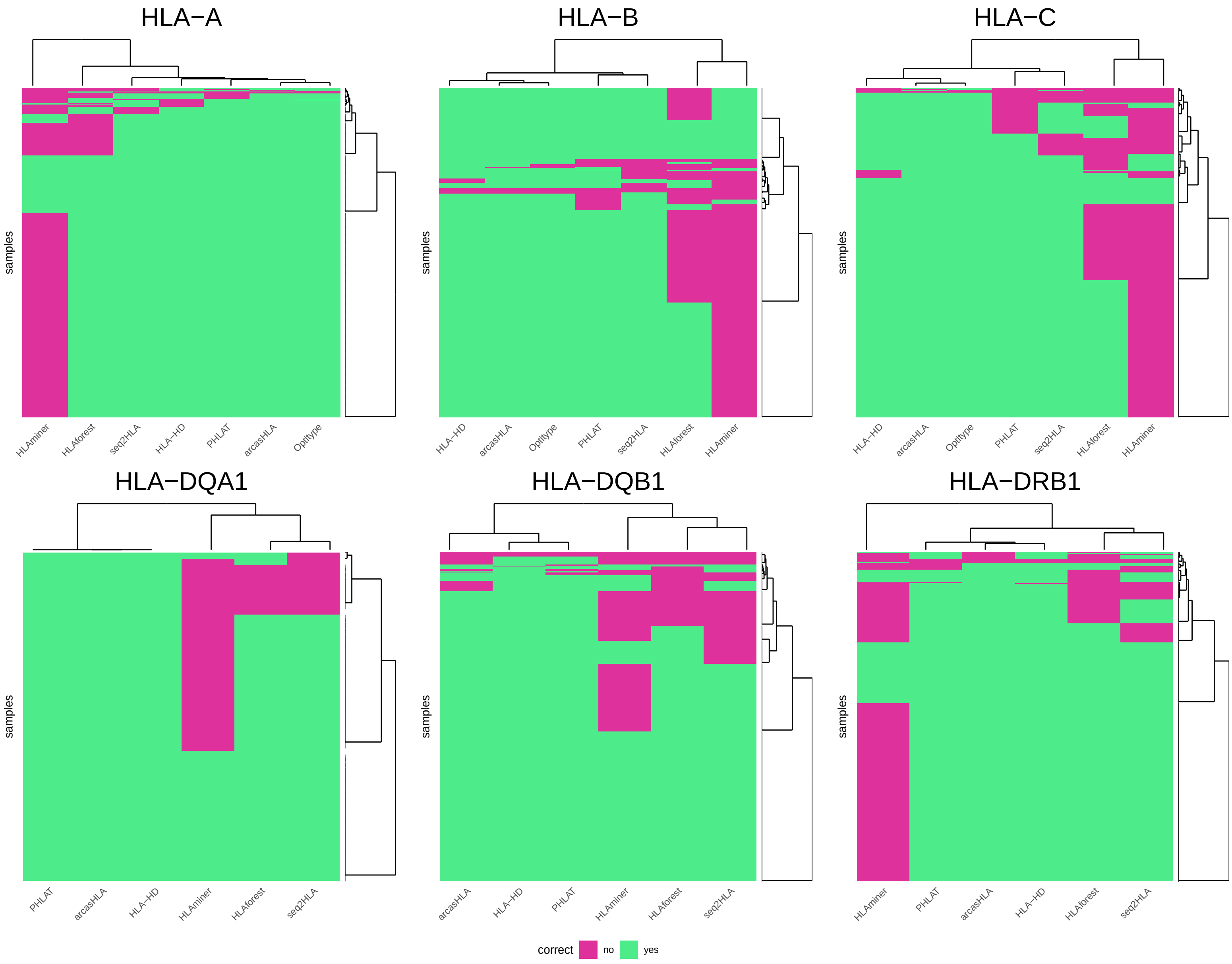


**Figure S10: Correctness of predictions on RNA data**
Heatmap indicating correctness of predictions on RNA data for each sample (rows) and tool (columns). Hierarchical clustering was applied on tools and samples. Dendrogram for the tools is shown on top of the plots. Dendrogram for the samples is shown right of the plots.

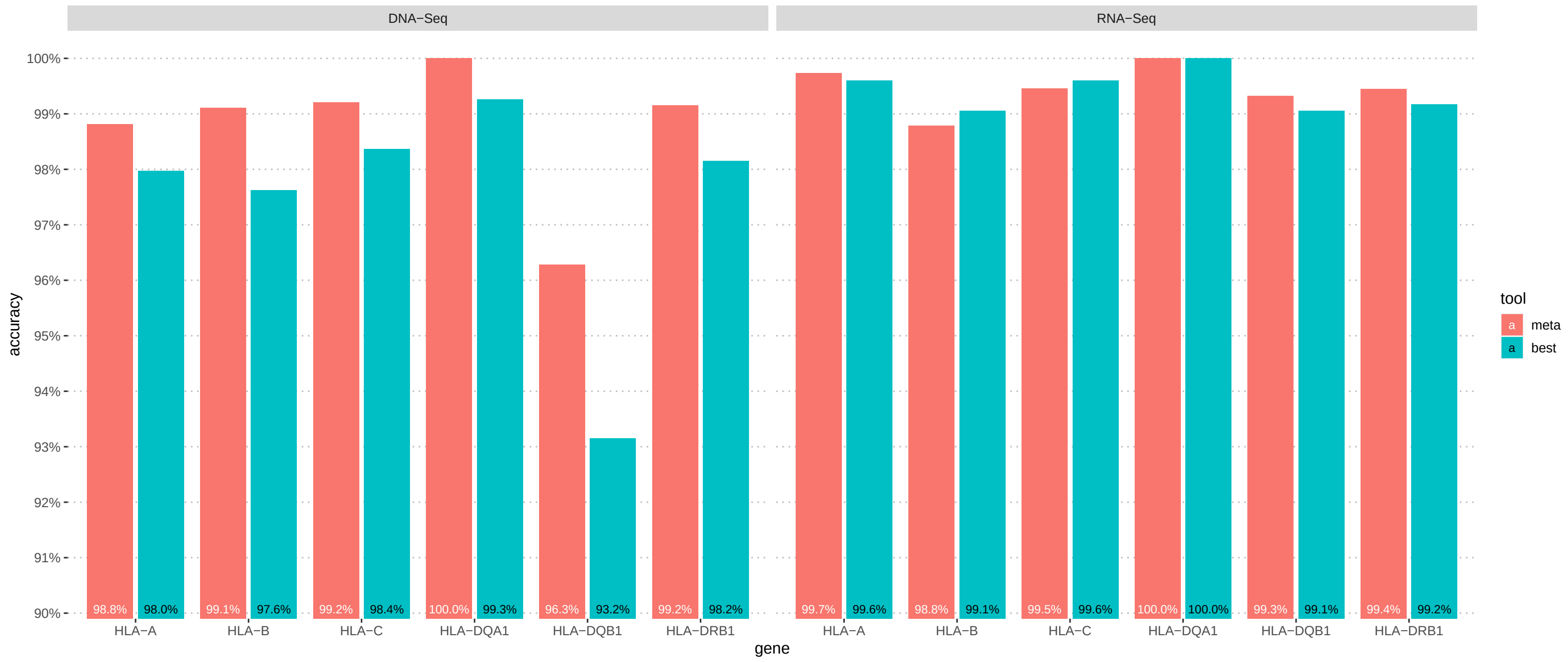


**Figure S11: Comparison of accuracies of all-tool metaclassifier with best performing individual tool per gene.**
Barplots comparing the accuracy of the best tool for each gene and data type to the accuracy of a classifier that chooses an HLA genotype from the output of all tools that support that data type and gene based on a majority voting rule. Bars in a red correspond to the accuracies of the voting classifier. Bars in blue correspond to the accuracies of the best individual tool for that gene.

### Supplementary tables

#### Table S1: Motivation for exclusion of 9 tools from the study

Overview of tools that were not benchmarked in our study and the reason for their exclusion. Excluded tools were either not *freely available for academic use*, do not use *FASTQ or BAM input files from WGS, WES and/or RNA-Seq* experiments (e.g., enrichment of the HLA region prior to sequencing is needed) or we were not able to get them *running on Ubuntu 20.04*.

#### Table S2: Algorithmic characteristics of the 13 selected tools

Table that compares the 13 selected HLA genotyping algorithms. Methods differ in the method to align reads to the HLA allele reference sequences (alignment method) and in the subsequent allele prioritization approach (allele pair prioritization). The column *tool-specific* steps describes additional particularities of the tools.

#### Table S3: Overview of AFN datasets used for expected allele frequencies

Overview of populations (studies) from the *Allele Frequency Net database* which were used to compile the list of expected HLA allele frequencies.

#### Table S4: Comparison of our results with 7 other independent benchmark studies

The allele prediction accuracies obtained in our benchmark on the 1000 genomes data are compared with 7 other independent benchmark papers. Each score was assigned a colour code: red under 50%, yellow between 50% and 80% and green above 80%. A grey box indicates that a tool was not evaluated for the corresponding gene in that study. The second row of the table indicates the gene or MHC class, where *class I + II* corresponds to the overall (combined) accuracy for MHC class I and class II. For the Lee, Kiyotani, Bauer and Yu studies the tools were evaluated on data from the 1000 genomes project. The studies by Liu, Yi and Chen used a different in house dataset. DNA data always refers to WES data.
