## Supplementary tables S1-S4 for "Benchmark of tools for *in silico* prediction of MHC class I and class II genotypes from NGS data": S1.pdf

| | Freely available for academic use | FASTQ or BAM input files from WGS, WES and/or RNA-Seq | Running on Ubuntu 20.04 |
|---|---|---|---|
| *ALPHLARD(-NT)* | ✗ | | |
| *ATHLATES* | | | ✗ |
| *HLAProfiler* | | | ✗ |
| *HLAreporter* | | | ✗ |
| *HLAssign* | | ✗ | ✗* |
| *OncoHLA* | ✗ | | |
| *PolyPheMe* | ✗ | | |
| *SNP2HLA* | | ✗ | |
| *SOAP-HLA* | | | ✗ |

* Latest version of HLAssign is a Windows GUI tool
