## Supplementary tables S1-S4 for "Benchmark of tools for *in silico* prediction of MHC class I and class II genotypes from NGS data": S2.pdf

| | Alignment method | Allele pair prioritization | | | Tool-specific steps |
|---|---|---|---|---|---|
| | | **independent scoring for 2 alleles** | **PHRED score used** | **prior population frequencies** | |
| **Kourami** | BWA-MEM | no | yes | no | Graph-guided assembly (post-alignment) |
| **HLA*LA** | BWA-MEM | no | yes | no | Graph-based optimization of alignment |
| **arcasHLA** | Kallisto (pseudo-alignment) | yes | no | yes | Iterative read re-allocation |
| **HLA-HD** | Bowtie2 | no | no | yes* | Separate mapping for introns and exons and for forward and reverse end reads |
| **PHLAT** | Bowtie2 | no | yes | yes* | Score calculated over and across SNP sites |
| **seq2HLA** | Bowtie2 | yes | no | no | Reads associated with first found allele removed before determining second |
| **xHLA** | DIAMOND | no | no | no | Iterative refinement |
| **Optitype** | RazerS3 | no | no | no | - |
| **Polysolver** | Novoalign | yes | yes | optional | - |
| **HLA-VBSeq** | BWA-MEM | yes | no | no | - |
| **HLAscan** | BWA-MEM | yes | no | no | Alleles discarded based on number of consecutive positions with no read aligned to it |
| **HLAforest** | Bowtie2 | yes | yes | no | Propagation of scores in alignment tree |
| **HLAminer** | BWA-backtrack | yes | no | no | - |

* only used for breaking-ties in case of ambiguities
