## Supplementary tables S1-S4 for "Benchmark of tools for *in silico* prediction of MHC class I and class II genotypes from NGS data": S3.docx

| Ethnic origin | Gene | AFN population ID | AFN population name |
| --- | --- | --- | --- |
| Black | HLA-A | 1480 | USA African American |
| Black | HLA-A | 1620 | USA African American Bethesda |
| Black | HLA-A | 2223 | USA African American pop 3 |
| Black | HLA-A | 2419 | USA African American pop 4 |
| Black | HLA-B | 1480 | USA African American |
| Black | HLA-B | 2223 | USA African American pop 3 |
| Black | HLA-B | 2419 | USA African American pop 4 |
| Black | HLA-C | 1480 | USA African American |
| Black | HLA-C | 2223 | USA African American pop 3 |
| Black | HLA-C | 2419 | USA African American pop 4 |
| Black | HLA-DPB1 | 2779 | USA African American pop 7 |
| Black | HLA-DQA1 | 1620 | USA African American Bethesda |
| Black | HLA-DQB1 | 2419 | USA African American pop 4 |
| Black | HLA-DQB1 | 2779 | USA African American pop 7 |
| Black | HLA-DRB1 | 1511 | USA Colorado Univ Cord Blood Bank African American |
| Black | HLA-DRB1 | 1620 | USA African American Bethesda |
| Black | HLA-DRB1 | 2223 | USA African American pop 3 |
| Black | HLA-DRB1 | 2419 | USA African American pop 4 |
| Black | HLA-DRB1 | 2779 | USA African American pop 7 |
| Caucasoid | HLA-A | 1359 | USA San Antonio Caucasian |
| Caucasoid | HLA-A | 1479 | USA Caucasian pop 2 |
| Caucasoid | HLA-A | 1619 | USA Caucasian Bethesda |
| Caucasoid | HLA-A | 2570 | USA Eastern European |
| Caucasoid | HLA-B | 1359 | USA San Antonio Caucasian |
| Caucasoid | HLA-B | 1479 | USA Caucasian pop 2 |
| Caucasoid | HLA-B | 1895 | USA Philadelphia Caucasian |
| Caucasoid | HLA-B | 2570 | USA Eastern European |
| Caucasoid | HLA-C | 1359 | USA San Antonio Caucasian |
| Caucasoid | HLA-C | 1479 | USA Caucasian pop 2 |
| Caucasoid | HLA-C | 1619 | USA Caucasian Bethesda |
| Caucasoid | HLA-C | 1895 | USA Philadelphia Caucasian |
| Caucasoid | HLA-DPA1 | 1279 | France Ceph |
| Caucasoid | HLA-DPA1 | 1401 | Spain Navarre Basques |
| Caucasoid | HLA-DPA1 | 2531 | Sweden pop 2 |
| Caucasoid | HLA-DPB1 | 2780 | USA Caucasian pop 5 |
| Caucasoid | HLA-DQA1 | 1619 | USA Caucasian Bethesda |
| Caucasoid | HLA-DQB1 | 1359 | USA San Antonio Caucasian |
| Caucasoid | HLA-DQB1 | 1895 | USA Philadelphia Caucasian |
| Caucasoid | HLA-DQB1 | 2780 | USA Caucasian pop 5 |
| Caucasoid | HLA-DRB1 | 1359 | USA San Antonio Caucasian |
| Caucasoid | HLA-DRB1 | 1513 | USA Colorado Univ Cord Blood Bank Caucasian |
| Caucasoid | HLA-DRB1 | 1586 | USA Caucasian Houston |
| Caucasoid | HLA-DRB1 | 1588 | USA Caucasian Pittsburgh |
| Caucasoid | HLA-DRB1 | 1619 | USA Caucasian Bethesda |
| Caucasoid | HLA-DRB1 | 1895 | USA Philadelphia Caucasian |
| Caucasoid | HLA-DRB1 | 2570 | USA Eastern European |
| Caucasoid | HLA-DRB1 | 2780 | USA Caucasian pop 5 |
